# Simulating alternative forest management in a changing climate on a *Pinus nigra* subsp. *laricio* plantation in Southern Italy

**DOI:** 10.1101/2022.05.12.491636

**Authors:** Riccardo Testolin, Daniela Dalmonech, Gina Marano, Maurizio Bagnara, Ettore D’Andrea, Giorgio Matteucci, Sergio Noce, Alessio Collalti

**Author notes:** contributed equally (Riccardo Testolin, Daniela Dalmonech).

## Abstract

Mediterranean pine plantations provide several ecosystem services but are particularly sensitive to climate change. Forest management practices might play a strategic role in the long-term adaptation of Mediterranean forests, but the joint effect of climate change and alternative management options in the near and far future have seldom been investigated together. Here, we developed a portfolio of management options and simulated the development of a Laricio pine (*Pinus nigra* subsp. *laricio*) stand in the Bonis watershed (southern Italy) from its establishment in 1958 up to 2095 using a state-of-the-science process-based forest model. The model was run under three climate change scenarios corresponding to increasing levels of atmospheric CO_2_ concentration, and seven management options with different goals, including post-disturbance management, wood production and renaturalization purposes. We analyzed the effect of climate change on annual carbon fluxes (i.e., gross and net primary production) and stocks (i.e., basal area and potential carbon woody stocks), as well as the impact of different management options compared to no management. Results show that, while climate change (i.e., warming and enriched atmospheric CO_2_ concentration) seems to increase carbon fluxes and stocks in the first half of the century, both show a substantial decrease in the second half, along with higher temperatures (+3 to +5 °C) and lower precipitation (−20% to −22%). When compared to no management, alternative options had a moderate effect on carbon fluxes over the whole simulation (between −6% and +7%) but overall carbon stocks were maximized by thinning interventions and the shelterwood system (+54% to +55%). We demonstrate that the choice of management exerts greater effects on the features of Laricio pine plantations than climate change alone. Therefore, silvicultural strategies might enhance potential stocks and improve forest conditions, with cascading positive effects on the provision of ecosystem services in Mediterranean pine plantations.

**Highlights:**

1. We simulated the development of a Laricio pine stand over 137 years under three different climatic scenarios and seven management options.
2. Carbon fluxes and stocks benefit from climate change (i.e., warming and enriched atmospheric CO_2_ concentration) in the first half of the century but show a marked decrease in the longer-term.
3. Forest management exerts a much stronger effect on these features than climate change alone.
4. Silvicultural options aimed at reducing stand density preserve and enhance carbon fluxes and stocks over the simulated time period.

## Introduction

Temperate forests play an important role in the Earth system Carbon (C) cycle by absorbing and storing a considerable amount of C in their aboveground and belowground compartments (Keith et al., 2009). Among these environments, Mediterranean forests account for 30% of the European forest cover and represent a net C-sink (FAO, 2018; Morán-Ordóñez et al., 2021). The Mediterranean basin is also a global biodiversity hotspot (Myers et al., 2000; Noce et al., 2016), with its forests harboring three times the number of tree species as the rest of Europe in a fourfold smaller area (Fady-Welterlen, 2005). These ecosystems play a key role in the livelihoods of local communities by providing food, timber, clean water, protection against soil erosion and micro-climatic regulation (Mazza et al., 2018; Morán-Ordóñez et al., 2021, 2020). At the same time, the Mediterranean basin is one of the main climate change hotspots on the planet (Diffenbaugh and Giorgi, 2012; Noce et al., 2017; Tuel and Eltahir, 2020). Indeed, the area is warming up 20% faster than the global average, precipitations are projected to decrease up to 20%, and extreme climatic events (e.g., heatwaves and droughts) are likely to increase both in frequency and intensity (D’Andrea et al., 2020; Lionello and Scarascia, 2018; Santini et al., 2014). These changing conditions could potentially reduce forest growth and prompt changes in forest dynamics (i.e., mortality and extensive dieback episodes) that, together with other disturbances, might limit the productivity and C-uptake capacity of Mediterranean forests (Gentilesca et al., 2017; Klein et al., 2019; Matteucci et al., 2013; Resco De Dios et al., 2007). By the end of this century, the cumulative effect of climate and land use change in the Mediterranean basin could trigger the transition from a positive (sink) to a negative (source) C-balance in the area, with inevitable and profound consequences on the persistence and dynamics of these ecosystems (Morales et al., 2007; Nolè et al., 2013; Pausas and Millán, 2019).

In this context, there is a high expectation towards the sustainable management of Mediterranean forests to counterbalance possible climate-change induced C-losses by preserving their sink and stock capability (Jandl et al., 2019; Reyer et al., 2015; Ruiz-Peinado et al., 2017; Vilà-Cabrera et al., 2018). Indeed, sustainable forest management practices can lower greenhouse gas emissions and contribute to climate change adaptation, while providing long-term livelihoods for communities by maintaining and enhancing ecosystem services (IPCC, 2019). This is especially critical for Mediterranean forests, as they have already undergone several millennia of human influence which resulted in the prevalence of mixed forest stands and conifer plantations (Ruiz-Benito et al., 2012). Among the latter, pine plantations were mainly established during the 20^th^ century to restore overexploited land, foster soil protection, and increase the production of existing forest stands, resulting in multiple forest restoration projects on a vast scale (Maestre and Cortina, 2004; Pausas et al., 2004). Despite the typical fast growing performances, Mediterranean pine plantations are particularly sensitive to the adverse effect of climate change and related disturbances (e.g., wildfires, drought, insect outbreaks; González-Sanchis et al., 2015; Martin-Benito et al., 2011; Navarro-Cerrillo et al., 2019; Resco De Dios et al., 2007; Ruiz-Benito et al., 2012), which might be further exacerbated by the lack of silvicultural treatments. This is particularly relevant in those mountainous areas characterized by limited accessibility and overall low economic revenue due to the high forest operation costs (Lerma-Arce et al., 2021; Proto et al., 2020). Therefore, management interventions in Mediterranean pine plantations aimed at promoting the progressive evolution of these stands towards more diverse and species-rich forests should be considered in order to ensure the future provision of ecosystem services in a changing climate (Nocentini et al., 2022).

Management strategies for climate change adaptation in Mediterranean forests are mainly translated into different thinning schemes – both in terms of intervention frequency and removal intensities – and ultimately through adjusted rotation periods (Resco De Dios et al., 2007). These adaptation measures (i) modulate C-stocks and C-uptake capacity, (ii) increase drought-stress resistance by reducing competition for water, and (iii) reduce losses of C use efficiency (net vs. gross primary production) by contrasting the aging of Mediterranean forests in the short-term, compared to the absence of management (del Río et al., 2017; González-Sanchis et al., 2015; Navarro-Cerrillo et al., 2019; Vilà-Cabrera et al., 2018). Despite the potential benefits of silvicultural practices aimed at enhancing the resilience of Mediterranean forests to future climate change impacts, the effects of management on the long-term forest adaptation are seldom investigated (Vilà-Cabrera et al., 2018), with the exception of few studies in high productivity regions (Manrique-Alba et al., 2020).

Process-based forest models provide a unique experimental framework to track the future responses of forest ecosystems to alternative management strategies under a changing climate (Gupta and Sharma, 2019; Keenan et al., 2011; Maréchaux et al., 2021; Reyer et al., 2015; Ruiz-Benito et al., 2020). Such models incorporate both empirical and mechanistic relations of the main ecophysiological processes which drive the response of forest stand development over decadal time periods (Gupta and Sharma, 2019; Keenan et al., 2011; Mäkelä et al., 2000) and can therefore help quantify the impacts of climate change and management on forest fluxes and stocks under changing environmental conditions. In an integrated scenario-analysis framework, process-based forest models can inform both the scientific and policy-oriented community of the forestry sector, thus supporting adaptation strategies in the Mediterranean basin (Keenan et al., 2011; Morán-Ordóñez et al., 2020; Vilà-Cabrera et al., 2018).

By means of a state-of-the-science process-based forest model (3D-CMCC-FEM; Three Dimensional - Coupled Model Carbon Cycle - Forest Ecosystem Model), we simulated the development of a Laricio pine stand in the Bonis experimental watershed (southern Italy) with the aim of providing insights on future adaptive management strategies of a Mediterranean pine plantation. We designed a wide portfolio of silvicultural strategies based on different forest management schemes which are currently applied in the study area and tested their effects on forest development under different climate change scenarios. Specifically, we aimed to 1) assess the impact of climate change alone on the forest C-budget including its annual productivity and stock capacity and, 2) evaluate the extent to which different silvicultural practices will affect C-balance up to the end of the 21^st^ century in one of the southernmost European forest sites.

## Materials and methods

### Study area and stand data collection

The Bonis experimental watershed is located in the mountain area of Sila Greca (39°28’49’’ N, 16°32’07’’ E; from 975 to 1330 m. a.s.l.) in the Calabria region, southern Italy, and represents one of the southernmost long-term experimental research sites in Europe. The catchment has a surface of 1.39 km^2^, a mean elevation of 1131 m a.s.l. and was firstly instrumented for hydrological monitoring in 1986. Almost 93% of the total area is covered by forests, dominated by ~60 years old Laricio pine stands, whose origin is mainly artificial (Callegari et al., 2003; Caloiero et al., 2017). The stands were planted in 1958 with an average density of 2425 saplings ha^-1^ (Nicolaci et al., 2015) and underwent a thinning treatment in 1993 which removed 25% of the basal area (BA) (Callegari et al., 2003). The climate is typically Mediterranean, with average annual precipitation of 915 mm and average temperature of 8.9 °C. The geological substrate is mainly composed of acid plutonic rocks and gravelly sands (Callegari et al., 2003). As part of the Euroflux-Carboitaly network, a tower for the measurement of eddy fluxes was installed in 2003 in a Laricio pine plantation within the study area (39°28’40’’ N, 16°32’05’’ E; Marino et al., 2005) and operated between 2005 and 2009. Furthermore, 14 circular 12 m-radius plots were established in 1993 before the thinning interventions and were resurveyed in 1999 and 2016. In each plot, for all trees with diameter at breast height (DBH; 1.3 m) > 2.5 cm, total height, crown insertion height and vitality were recorded (Collalti et al., 2017). The plot data have been used to parameterize and, together with the eddy fluxes data, to validate the model.

### Vegetation model and species parameterization

The 3D-CMCC-FEM forest model (v.5.6 BGC) is a biogeochemical, biophysical, and physiological process-based forest model developed to predict C, energy, and water fluxes coupled with stand development processes that determine relative stock changes in forest ecosystems (Collalti et al., 2019; Dalmonech et al., 2022). The model is designed to simulate the main physiological and hydrological processes at daily, monthly, and annual scales and at the species-specific level. The model requires data on initial forest stand conditions, including species composition, average tree DBH, height, stand age and tree density (number of trees per hectare). Both structural and non-structural tree C-pools are initialized at the beginning of the simulation and updated daily, monthly, or annually, depending on the processes. Furthermore, the model allows the simulation of different management scenarios by defining the intensity and the interval of removals, as well as the length of rotation periods and artificial replanting schemes, which can be varied through the simulation time. For a full description of key model principles and theoretical framework see also Collalti et al. (2020, 2019, 2018, 2016, 2014), Dalmonech et al. (2022), Engel et al. (2021), and Marconi et al. (2017).

The model was parameterized to simulate the development of a Laricio pine stand based on published literature (Lapa et al., 2017; Lebourgeois et al., 1998; Patenaude et al., 2008). When published information on the species was unavailable for a given ecophysiological parameter, we used the values reported for ecologically-close species following this order: other subspecies of *Pinus nigra* (Grossoni, 2014; Margolis et al., 1995; Móricz et al., 2018; Navarro-Cerrillo et al., 2016; Van Haverbeke, 1990), *Pinus pinaster* (Chiesi et al., 2007; Delzon et al., 2004; Mollicone et al., 2002), *Pinus sylvestris* (Collalti et al., 2019; Yuste et al., 2005) or, more generally and in few cases, other evergreen species (Arora and Boer, 2005; Dewar et al., 1994; Poulter et al., 2010). All parameter values and sources are reported in Supplementary Information Table S1.

### Climate and atmospheric CO_2_ data

The 3D-CMCC-FEM requires as climatic inputs daily values of solar radiation (MJ m^−2^), temperature (°C), precipitation (mm) and vapor pressure deficit (hPa). Such data, from 1958 to 2016, were derived for the Bonis watershed using the mountain microclimate simulation model MT-CLIM (Thornton and Running, 1999) forced by temperature and precipitation series measured by the nearby Cecita meteorological station (39°23’51’’ N, 16°33’24’’ E; 1180 m a.s.l.). This dataset was used to perform historical simulations for model validation.

To simulate the development of the Laricio pine stand up to the end of the 21^st^ century, we employed a set of climate data covering the 1976 - 2095 period at 0.0715° spatial resolution (~8 km) (Bucchignani et al., 2016; Zollo et al., 2016). This highly resolved climate data are based on the regional climate model COSMO-CLM (Rockel et al., 2008) driven by the CMCC-CM global model (Scoccimarro et al., 2011) using the 20C3M forcing (i.e., observed emissions) for the period 1976 - 2005, and two IPCC emission scenarios from 2006 onwards: the intermediate emission scenario RCP4.5 and the high emission scenario RCP8.5 (Moss et al., 2010; van Vuuren et al., 2011). The RCP4.5 scenario assumes that the total radiative forcing is stabilized, shortly after 2100, to 4.5 Wm^−2^ (approximately 650 ppmv CO_2_-equivalent) by employing various technologies and strategies to reduce greenhouse gas emissions. The RCP8.5 is characterized by increasing emissions and high greenhouse gas concentration levels, leading to 8.5 Wm^-2^ in 2100 (approximately 1370 ppmv CO_2_-equivalent). Modeled temperature and precipitation data were bias corrected following the approach adopted and described in Sperna Weiland et al. (2010), starting from the observed series of the same variables. As an observational dataset for the bias correction the downscaled daily E-OBS dataset (v 10.0) at 1 km resolution (Maselli et al., 2012) was used. Additionally, we simulated a no climate change (NOCC) dataset as a benchmark scenario for the period 2006 - 2095 by randomly sampling each day in sequence from the bias-corrected COSMO-CLM dataset between 1990 and 2005. As the COSMO-CLM data were only available starting from 1976, we used the MT-CLIM climatic dataset described above for the 1958 - 1975 period.

Measured values of global annual atmospheric CO_2_ concentration (ppmv) were derived from Meinshausen et al. (2011), while values consistent to the abovementioned emission scenarios were provided by Dlugokencky and Tans (2014). The atmospheric CO_2_ concentrations for the NOCC scenario were simulated by randomly sampling each year in sequence between 1990 and 2005 from Meinshausen et al. (2011).

To assess the departure of projected climate change from the baseline NOCC scenario, we calculated the mean relative change in temperature, precipitation, vapor pressure deficit and atmospheric CO_2_ concentration for the two RCP scenarios within two different time windows: near future (NF; 2025 - 2055) and far future (FF; 2065 - 2095). 95% confidence intervals were estimated as ± 1.96 times the standard error. Disjoint confidence intervals were considered as a conservative indication of statistically significant differences among scenarios.

### Model evaluation

Model performances were evaluated by simulating the development of a representative Laricio pine stand in the Bonis watershed from its establishment in 1958 to the last field measurements occurred in 2016, which includes the thinning in 1993. The model was initialized in 1958 with an initial density of 2425 saplings per hectare (DBH: 1 cm, height: 1.3 m, age: 4 years; Nicolaci et al., 2015), considering the average elevation of the watershed (1131 m.a.s.l.), the average soil texture (clay: 20%; silt: 26%; sand: 54%) and depth (100 cm) (Buttafuoco et al., 2005; Moresi et al., 2020). The evaluation was carried out by comparing the resulting simulated mean annual DBH and tree density to the values measured at the field plots in 1993 (before thinning), 1999 and 2016, as well as to the estimations provided by Callegari et al. (2003) for low and high density Laricio pine plantations in the Bonis watershed for 1986, 1993 (before and after thinning) and 1999. Additionally, a micrometeorological validation of daily gross primary productivity (GPP) was carried out by comparing the simulated values to those obtained by the eddy covariance tower. Only the measurements up to 2008 were considered, as the 2009 dataset presented major gaps in the daily time series. Among the selected data, we excluded all days with a quality control flag lower than 0.6 which were then removed from the simulation settings as described in Collalti et al. (2018). The comparisons were carried out for each year, as well as for the daily averages of the two years, by calculating root mean squared error (RMSE), coefficient of determination (R^2^) and modeling efficiency (ME). The latter index provides information about modeling performance on a relative scale: ME = 1 indicates a perfect fit, ME = 0 reveals that the model is no better than a simple average, while negative values indicate poor performance (Bagnara et al., 2015; Vanclay and Skovsgaard, 1997).

### Forest management scenarios

For each of the three climate scenarios (i.e., NOCC, RCP4.5, RCP8.5) we simulated forest management by mimicking seven different silvicultural options reflecting different goals (Table 1), resulting in a total of 21 different model runs. All the options were simulated to take place after 2016, i.e., the last year of field measurements. The scenarios cover several management objectives including post-disturbance management, wood production and renaturalization and reflect the state-of-the-science of management options applied to this region of the Italian Apennines (Cantiani et al., 2018). The first option (‘*no management*’) represents the natural development of the forest left without human intervention, while the second option (‘*natural regeneration*’) reproduces natural forest regeneration following a major disturbance event (e.g., wildfire), simulated as a clear-cut after 80 years from planting (i.e., around the time when atmospheric aridity start increasing while the fuel load is still high). The regeneration is simulated as a prescribed replanting, with density of saplings derived from the estimated tree density of natural Laricio pine stands in 1986 (Callegari et al., 2003) by going backwards to 1958 assuming a 1% annual mortality rate (Andrus et al., 2021). Two options simulating different thinning intensities – ‘*light*’ and ‘*heavy*’, corresponding to a 28% and 35.5% reduction of BA, respectively – at an interval of 15 years are proposed in order to reproduce silvicultural interventions aimed at favoring natural forest dynamics. Indeed, at intermediate stages of stand development, pine forests can benefit from thinnings aimed specifically at improving their degree of stability (Cantiani et al., 2005; Cantiani and Piovosi, 2008). Selective thinnings induce an increase in mechanical stability, favor structural diversity, and reduce inter-tree competition for water, light, and nutrients (del Río et al., 2017; Marchi et al., 2018). However, tending and thinning interventions still represent a major passive management item in terms of costs and are often avoided in public forests resulting in a progressive degeneration of stand structure (Ahtikoski et al., 2021; Niskanen and Väyrynen, 2001). An additional, production-oriented option (‘*patch clearcut*’) simulating a complete harvest followed by replanting 80 years after the establishment of the plantation is also included. Yet, the shelterwood system represents a more sustainable alternative to clear-cutting and patch cuttings by ensuring a progressive and constant light availability to the forest floor. The practice favors regeneration while modulating the competition for light and water resources with herbs and shrubs (not considered here), and allows higher revenues (Brichta et al., 2020; Cantiani et al., 2018; Montoro Girona et al., 2018). Therefore, we simulated two shelterwood options: ‘*shelterwood A*’, consisting of two light thinnings (20% reduction of BA) with a 10 year interval, followed by an establishment cut after 80 years from the original planting (80% reduction of BA) and a removal cut 10 year later; ‘*shelterwood B*’, defined by a delayed establishment cut after 90 years, preceded by three heavier thinnings (28.5% reduction of BA) and followed by a removal cut after 10 years. In both cases, the establishment cut is followed by natural regeneration of the same species.

**Table 1.** Summary of simulated management options. Abbreviations: r = rotation period; thBA = basal area removed with thinning; thINT = time interval between thinnings.

| Option | Detail | Objective | r | thBA | thINT | replanting | Description |
| --- | --- | --- | --- | --- | --- | --- | --- |
|  |  |  | year | % | year | n saplings ha<sup>-1</sup> |  |
| **No management** | No interventions | - | - | - | - | - | This option simulates only the documented thinning in 1993 (25% of BA). |
| **Natural regeneration** | Clearcut + natural regeneration | Post disturbance (wildfire) | 80 | - | - | 5013 | Clear-cut after 80 years from plantation establishment (year: 2038). After that, natural regeneration follows. |
| **Light thinning** | Multiple thinning interventions | Biodiversity / Renaturalization | - | 28 | 15 | - | 4 light thinnings (years: 2017, 2032, 2047, 2062). |
| **Heavy thinning** | Multiple thinning interventions | Biodiversity / Renaturalization | - | 35.5 | 15 | - | 4 heavy thinnings (years: 2017, 2032, 2047, 2062). |
| **Patch clearcut** | Clearcut + artificial regeneration (replanting) | Production / Commercial forest | 80 | - | - | 2425 | Complete harvest after 80 years from plantation establishment (year: 2038). After that, the same number of trees as in 1958 is replanted. |
| **Shelterwood A** | Thinnings | Production / Commercial forest | - | 20 | 10 | - | 2 light thinnings (years: 2017, 2027), 1 heavy thinning (establishment cut) in 2038 followed by natural regeneration, harvest (removal cut) in 2048. |
|  | Establishment cut |  | 80 | 80 | - | 5013 |  |
|  | Removal cut |  | 90 | 100 | - | - |  |
| **Shelterwood B** | Thinnings | Production / Commercial forest | - | 28.5 | 10 | - | 3 light thinnings (years: 2017, 2027, 2037), 1 heavy thinning (establishment cut) in 2048 followed by natural regeneration, harvest (removal cut) in 2058. |
|  | Establishment cut |  | 90 | 80 | - | 5013 |  |
|  | Removal cut |  | 100 | 100 | - | - |  |

### Analysis of simulation outputs

To assess the impacts of climate change and management on stand structure and function, we evaluated the temporal trends of GPP, net primary productivity (NPP), potential C-woody stocks (pCWS; i.e., the sum of standing woody biomass and harvested woody products when no decay is assumed) and BA. We chose these variables among all model outputs as they are key components of the forest C-budget and forest structure, representing the physiologically and structurally inherent capacity of trees to sequester and stock atmospheric CO_2_ on the short- (i.e., GPP and NPP) to long-term (i.e., pCWS and BA). At the same time, these outputs are key variables relevant to decision makers to assess stand growth changes and current standing biomass, as well as to make appropriate management decisions. Notably, we considered pCWS as representative of the maximum attainable C-stock capacity to quantify the inherent capability of trees to sequester and store C over medium-to long-time periods. Only in the ‘*natural regeneration*’ option, we assumed the destruction of the stocks following a forest fire.

We analyzed the overall effect of climate change by calculating the mean relative change of the abovementioned outputs between the RCP and the NOCC scenarios within the NF and FF time windows. The results were then averaged across all seven management options. Similarly, to assess the effect of management, we calculated the mean values of the target outputs for each option, as well as the relative change between each management option and no management, here considered as the baseline, averaging the outputs of the three climate scenarios. Apart from the NF and FF time windows, these results were also provided for the whole simulation starting from 2006 (i.e., the starting year of the climatic scenarios; ALL time window). To visualize the whole time series, we performed a *loess* fit of the simulated outputs for each management option with a span of 0.5 to reduce noise from interannual variability. Climate scenarios were considered jointly, thus representing the interval of values between the absence of climate change and the worst case scenario. 95% confidence intervals of each mean relative change were estimated and used to identify significant differences as described above. All data visualization and analyses were performed with R (R Core Team, 2021).

## Results

### Model evaluation

The simulated mean stand DBH of Laricio pine plantations in the Bonis watershed was 18.1 cm in 1986, 20.5 cm in 1993 before the thinning, 21 cm in 1993 after the thinning, and 24.3 cm in 1999. In the same years, Callegari et al. (2003) reported a mean stand DBH range of 18 - 20.2 cm, 19.8 - 21.8 cm, 20.8 - 22.8 cm and 23.8 - 27.4 cm, respectively, for high and low density plantations. At the forest plots, a mean stand DBH of 22.2 ± 2.4 cm was estimated in 1993 before the thinning, which increased to 25.9 ± 3.7 cm in 1999 and to 33.7 ± 3.3 cm in 2016. The simulated value for in 2016, was 33.6 cm (Table 2; Figure 1a). As for tree density, the model simulated 1620 trees ha^−1^ in 1986, 1276 trees ha^−1^ in 1993 before the thinning, 948 trees ha^−1^ in 1993 after the thinning, 894 trees ha^−1^ in 1999 and 474 trees ha^−1^ in 2016. The values measured at the forest plots were 1491 ± 382 trees ha^−1^, 975 ± 376 trees ha^−1^ and 522 ± 231 trees ha^−1^ in 1993 before the thinning, 1999 and 2016, respectively. Similarly, Callegari et al. (2003) reported a range of 1250 - 2200 trees ha^−1^, 1162 - 1701 trees ha^−1^, 800 - 1150 trees ha^−1^ and 775 - 1102 trees ha^−1^ in 1986, 1993 before thinning, 1993 after thinning and 1999, respectively (Table 2; Figure 1b).

**Figure 1.**
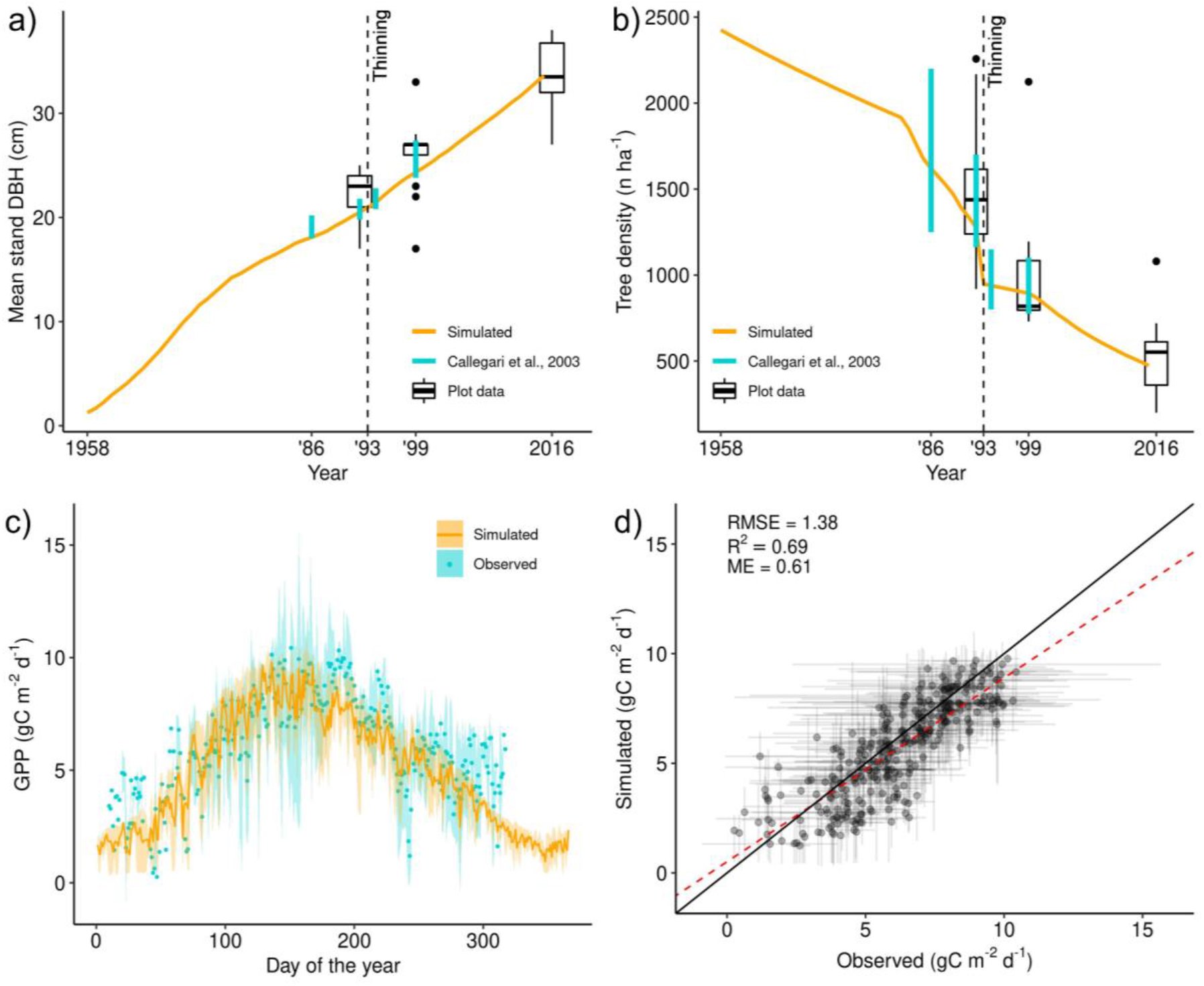
Evaluation of (a) simulated mean stand DBH and (b) tree density against the values reported by Callegari et al. (2003) and measured within the sampling plots. Evaluation of the average simulated daily GPP against the values obtained by the eddy covariance tower at the Bonis watershed in the years 2005 - 2009 (c, d). The solid line represents the mean simulated value. The points represent the mean values derived by eddy covariance measurements. Shaded areas (c) and error bars (d) are the interval between the minimum and maximum values for a given day.

**Table 2.** Simulated values of mean stand DBH and tree density (in bold) against those reported by Callegari et al. 2003 (range between low and high density plantations) and measured at the sampling plots (mean and standard deviation). The reported simulated values for 1993 (before thinning) and 1993 (after thinning) are for the years 1992 and 1993, respectively.

|  | **1986** | **1993**<br>**(before thinning)** | **1993**<br>**(after thinning)** | **1999** | **2016** |
| --- | --- | --- | --- | --- | --- |
| *Mean stand DBH (cm)* |  |  |  |  |  |
| **Simulated** | **18.1** | **20.5** | **21** | **24.3** | **33.6** |
| **Callegari et al. 2003** | 18 - 20.2 | 19.8 - 21.8 | 20.8 - 22.8 | 23.8 - 27.4 | - |
| **Plot data** | - | 22.2 ± 2.4 | - | 25.9 ± 3.7 | 33.7 ± 3.3 |
| *Tree density (n trees ha<sup>-1</sup>)* |  |  |  |  |  |
| **Simulated** | **1620** | **1276** | **948** | **894** | **474** |
| **Callegari et al. 2003** | 1250 - 2200 | 1162 - 1701 | 800 - 1150 | 775 - 1102 | - |
| **Plot data** | - | 1491 ± 382 | - | 975 ± 376 | 522 ± 231 |

Goodness-of-fit metrics of the four-year average trend of simulated daily GPP against values derived by the eddy covariance tower were RMSE = 1.38 gC m^−2^ d^−1^, R^2^ = 0.69 and ME = 0.6 (Figure 1c,d). As for the daily GPP of each year, the model reproduced the annual trends, albeit with different accuracy (Figure S1).

### Climate change scenarios

On average, atmospheric CO_2_ concentration increased to 461 - 494 ppmv in NF and to 530 - 761 ppmv in FF, according to the RCP4.5 and RCP8.5 scenarios, respectively. At the same time, mean temperatures at the Bonis watershed under the RCP4.5 scenario are projected to increase by 1.2 °C (9%) in NF and 3 °C (23%) in FF, compared to NOCC. According to the RCP8.5 scenario, the increase will be by 1.8 °C (14%) and 5 °C (39%). Vapor pressure deficit will also increase by 13% in NF and 31% in FF under the RCP4.5 scenario compared to NOCC, while the increase will be by 18% and 59% under the RCP8.5 scenario. No significant change in precipitation is predicted in NF for both scenarios, while a reduction of 20% and 22% is predicted in FF, respectively for the RCP4.5 and RCP8.5 scenarios, compared to NOCC (Table S2; Figure S2).

According to the RCP4.5 scenario, GPP will increase by 4.7% in NF and by 3% in FF, compared to NOCC. The increase according to the RCP8.5 scenario will be 7.8% in NF and 1.9% in FF. pCWS will not change in NF and will decrease by 1.8% in FF under RCP4.5, while it will slightly increase in NF (0.7%) and decrease in FF (−2.2%) under RCP8.5, compared to NOCC. BA will increase under both scenarios in NF (1.5% for RCP4.5 and 2.3% for RCP8.5) and decrease in FF (−1.2% for RCP4.5 and −1.9% for RCP8.5). No significant differences in NPP were detected for NF while a 11.7% reduction will take place in FF under the RCP4.5 scenario, and an even stronger 23% decrease is projected under the RCP8.5 scenario (Figure 2; Table S3).

**Figure 2.**
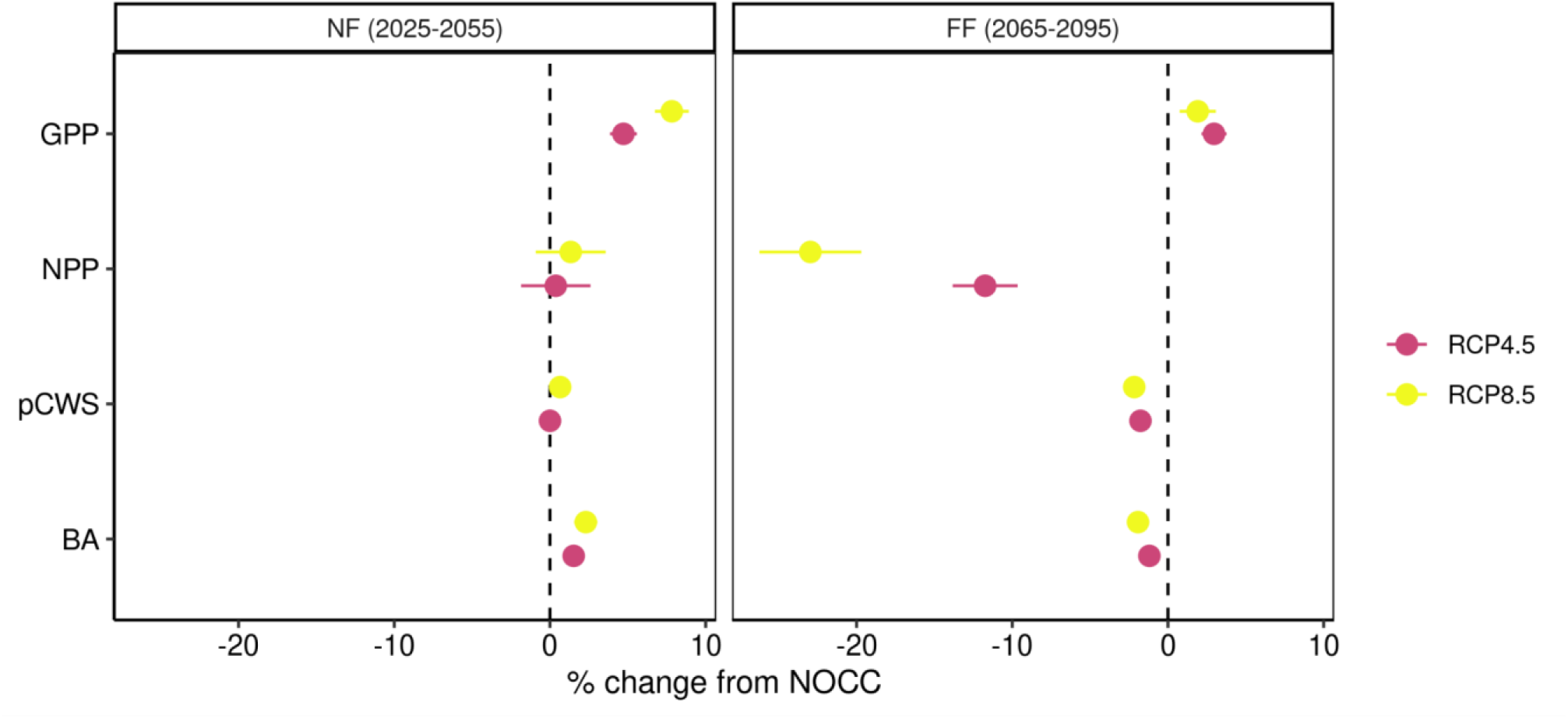
Relative change of simulation outputs between RCP4.5 and RCP8.5 climatic scenarios compared to the baseline NOCC scenario within the NF and FF time windows. The percentages were averaged across all seven management options. The error bars are the 95% confidence intervals.

### Forest management scenarios

Within the NF time window, the simulation under the ‘*no management*’ option exhibited the highest mean values for GPP (1636 gC m^−2^ y^−1^), NPP (559 gC m^−-2^ y^−1^) and BA (42 m^2^ ha^−1^), while the ‘*patch clearcut*’ scenario showed the lowest values of the same variables (GPP: 1221 gC m^−2^ y^−1^; NPP: 453 gC m^−2^ y^−1^; BA: 24 m^2^ ha^−1^). As for pCWS, the highest mean values were exhibited by the ‘*shelterwood B*’ option (168 tC ha^−1^), while the lowest were found in the ‘*natural regeneration*’ option (64 tC ha^−1^). The ‘*shelterwood A*’, ‘*shelterwood B*’, ‘*patch clearcut*’ and ‘*natural regeneration*’ options exhibited a similar decrease in GPP (between −17% and −25%) and BA (between −30% and −41%) compared to ‘*no management*’, while the ‘*light*’ and ‘*heavy thinning*’ options presented a similarly lower decrease (−4% and −7% for GPP; −11% and −16% for BA, respectively). As for NPP, ‘*light*’ and ‘*heavy thinning*’ showed a decrease of 2% and 3%, while ‘*natural regeneration*’ and ‘*patch clearcut*’ presented the greatest decrease (−14% and −18%); ‘*shelterwood A*’ and ‘*shelterwood B*’ exhibited intermediate values at −6% and −9% of NPP compared to ‘*no management*’ option. Increases in pCWS were between 37% and 46% for thinning and shelterwood options, while the ‘*patch clearcut*’ option exhibited a 4% increase compared to ‘*no management*’. The ‘*natural regeneration*’ option showed a 42% decrease (Table 3; Figure 3 and 4; Figure S3).

**Figure 3.**
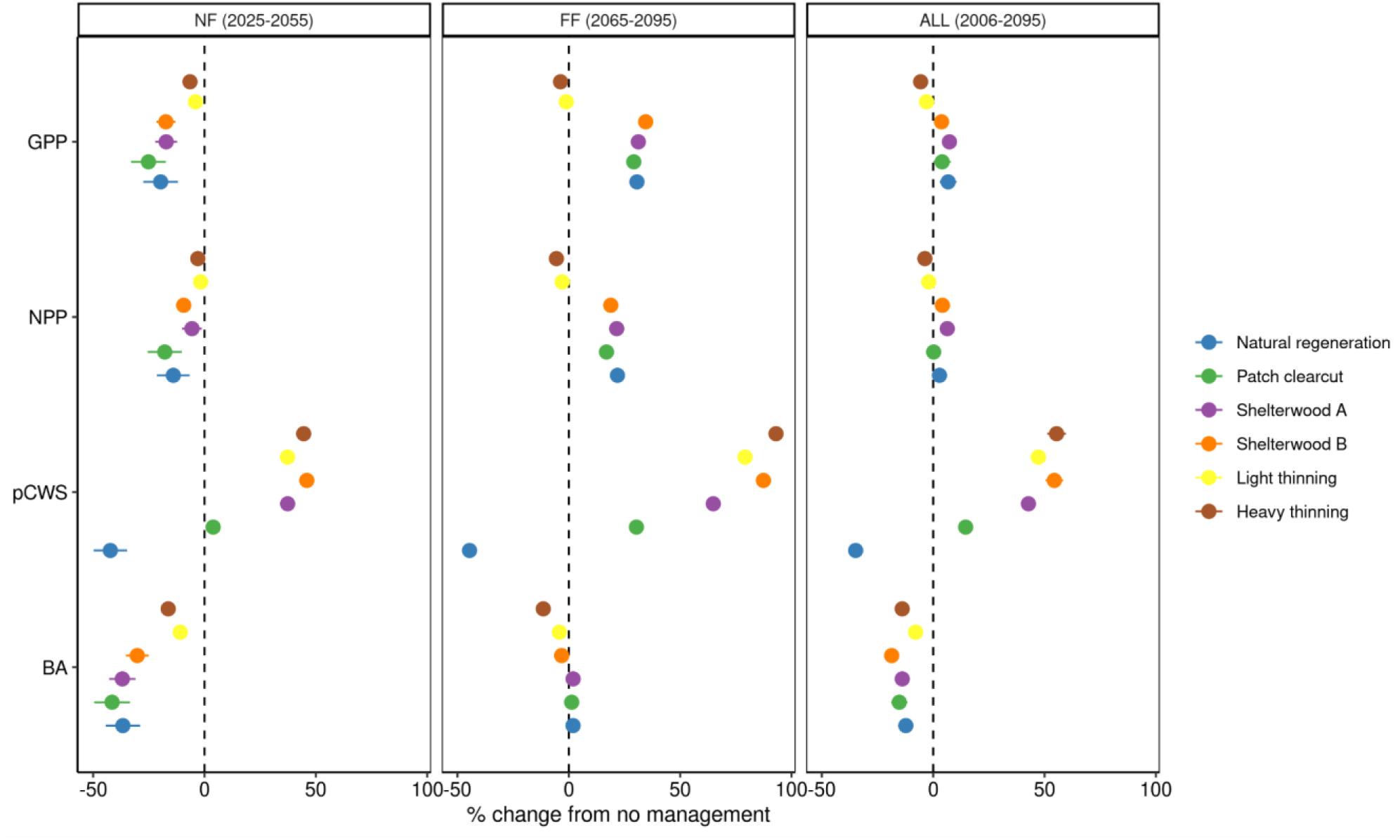
Relative change of modeled outputs according to different management options compared to the baseline ‘*no management*’ scenario within the NF, FF and ALL time windows. The error bars are the 95% confidence intervals.

**Figure 4.**
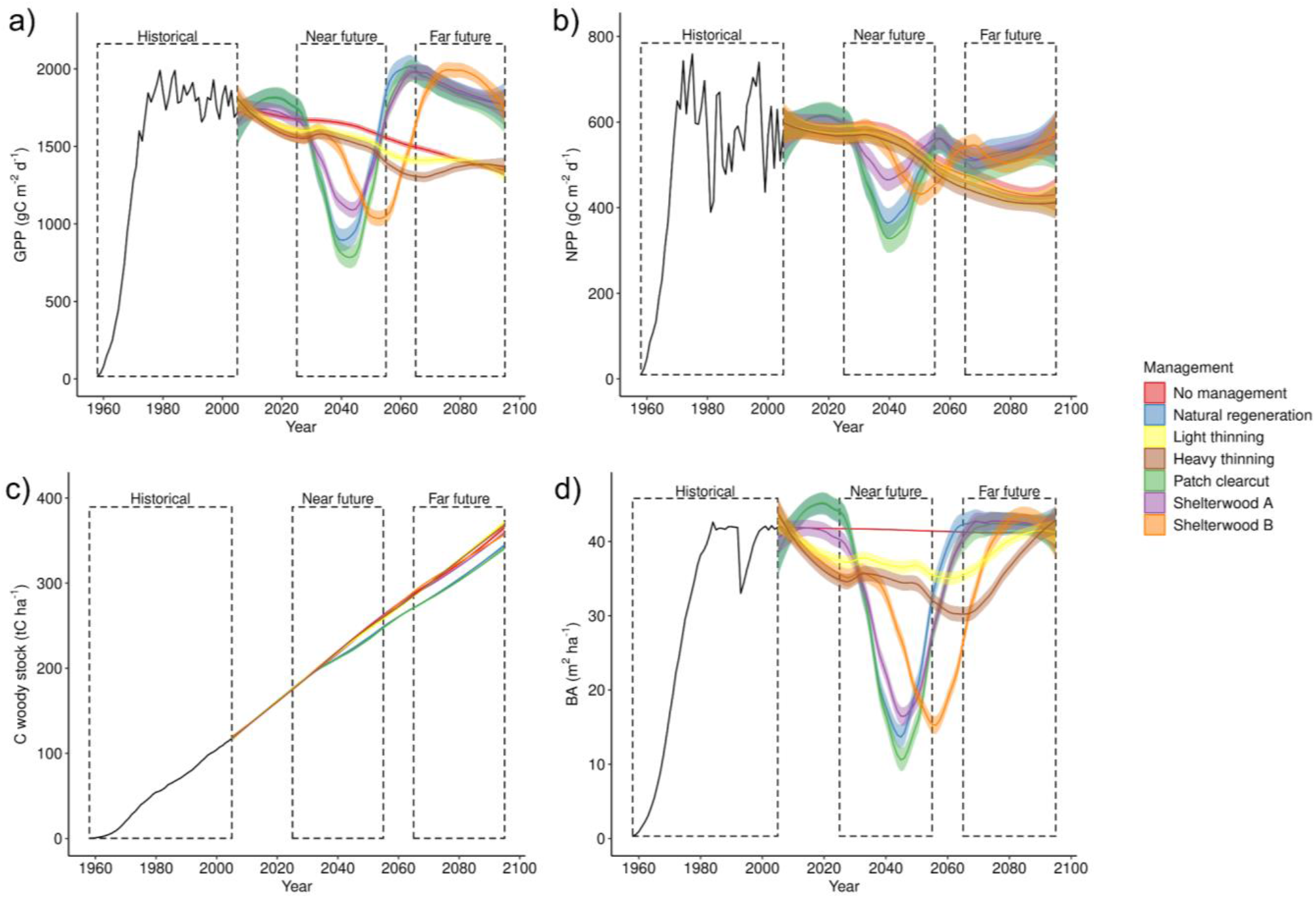
Simulated GPP (a), NPP (b), pCWS (c) and BA (d) according to seven management options. Black lines are the historical simulations from 1958 to 2005. Solid lines from 2006 onwards are a *loess* fit of the outputs produced by different climate scenarios (NOCC, RCP4.5, RCP8.5) for each management option. Shaded areas are the confidence intervals of the fit and represent the climatic variability among scenarios within each management option.

**Table 3.** Mean values of selected model outputs for seven management options and three time windows. The values have been averaged across all climate scenarios. Relative changes between each option and the baseline ‘*no management*’ scenario are reported in brackets. The highest and lowest values when compared to the ‘*no management*’ scenario are reported in bold.

|  | Near future (2025 - 2055) |  |  |  | Far future (2065 - 2095) |  |  |  | All (2006 - 2095) |  |  |  |
| --- | --- | --- | --- | --- | --- | --- | --- | --- | --- | --- | --- | --- |
|  | GPP<br>(gC m<sup>-2</sup> y<sup>-1</sup>) | NPP<br>(gC m<sup>-2</sup> y<sup>-1</sup>) | pCWS<br>(tC ha<sup>-1</sup>) | BA<br>(m<sup>2</sup> ha<sup>-1</sup>) | GPP<br>(gC m<sup>-2</sup> y<sup>-1</sup>) | NPP<br>(gC m<sup>-2</sup> y<sup>-1</sup>) | pCWS<br>(tC ha<sup>-1</sup>) | BA<br>(m<sup>2</sup> ha<sup>-1</sup>) | GPP<br>(gC m<sup>-2</sup> y<sup>-1</sup>) | NPP<br>(gC m<sup>-2</sup> y<sup>-1</sup>) | pCWS<br>(tC ha<sup>-1</sup>) | BA<br>(m<sup>2</sup> ha<sup>-1</sup>) |
| **No management (baseline)** | 1636 | 559 | 115 | 42 | 1415 | 443 | 139 | 41 | 1566 | 518 | 121 | 41 |
| **Natural regeneration** | 1309 (-20) | 475 (-14) | **64 (-42)** | 26 (-37) | 1846 (31) | **536 (22)** | **77 (-45)** | **42 (2)** | 1647 (7) | 522 (3) | **76 (-35)** | 36 (-12) |
| **Light thinning** | **1569 (-4)** | **549 (-2)** | 158 (37) | **37 (-11)** | 1397 (-1) | 430 (-3) | 250 (79) | 39 (-4) | 1518 (-3) | 508 (-2) | 183 (47) | **38 (-8)** |
| **Heavy thinning** | 1530 (-7) | 542 (-3) | 166 (45) | 35 (-16) | **1359 (-4)** | **419 (-6)** | **269 (93)** | **37 (-11)** | **1477 (-6)** | **500 (-4)** | **193 (55)** | 36 (-14) |
| **Patch clearcut** | **1221 (-25)** | **453 (-18)** | 119 (4) | **24 (-41)** | 1827 (29) | 515 (17) | 181 (30) | 42 (1) | 1605 (4) | 509 (0) | 141 (15) | 35 (-15) |
| **Shelterwood A** | 1352 (-17) | 522 (-6) | 158 (37) | 26 (-37) | 1856 (31) | 534 (21) | 229 (65) | 42 (2) | **1657 (7)** | **541 (6)** | 176 (43) | 36 (-14) |
| **Shelterwood B** | 1356 (-17) | 507 (-9) | **168 (46)** | 29 (-30) | **1901 (34)** | 524 (19) | 261 (87) | 40 (-3) | 1603 (4) | 532 (4) | 192 (54) | **34 (-19)** |

As for the FF time window, mean GPP was the highest under the ‘*shelterwood B*’ option (1901 gC m^−2^ y^−1^), while mean NPP was the highest under the ‘*natural regeneration*’ option (536 gC m^−2^ y^−1^). Mean pCWS was maximized with ‘*heavy thinning*’ (269 tC ha^−1^), while the highest simulated BA was tied between the ‘*natural regeneration*’ and ‘*shelterwood A*’ options (42 m^2^ ha^−1^). The ‘*heavy thinning*’ option led to the lowest mean GPP (1359 gC m^−2^ y^−1^), NPP (419 gC m^−2^ y^−1^) and BA (37 m^2^ ha^−1^), while the lowest mean pCWS emerged under the ‘*natural regeneration*’ simulation (77 tC ha^−1^) (Table 3; Figure 3). Overall, ‘*natural regeneration*’, ‘*patch clearcut*’, ‘*shelterwood A*’ and ‘*shelterwood B*’ options exhibited a similar increase in GPP (between 29% and 34%), NPP (between 17% and 22%) and BA (between 1% and 3%), compared to ‘*no management*’. Conversely, ‘*light*’ and ‘*heavy thinning*’ showed a decrease in GPP (−1% and −4%), NPP (−3% and −6%) and BA (−4% and −11%). pCWS increased between 79% and 93% under the thinning and shelterwood options, while it showed a 30% increase with ‘*patch clearcut*’ and a 45% decrease under the ‘*natural regeneration*’ option (Table 3; Figure 3 and 4; Figure S3).

Between 2006 and 2095, GPP was maximized under the *‘natural regeneration’, ‘patch clearcut’, ‘shelterwood A’* and ‘*shelterwood B*’ options (1603 - 1657 gC m^−2^ y^−1^), corresponding to a 4% to 7% increase compared to ‘*no management*’ (1566 gC m^−2^ y^−1^), while the thinning options showed the lowest values (1477 - 1518 gC m^−2^ y^−1^) and a decrease between 3% and 6%. NPP showed a similar trend, with the *‘natural regeneration’, ‘shelterwood A’* and ‘*shelterwood B*’ options exhibiting the highest values (522 - 541 gC m^−2^ y^−1^), corresponding to an increase between 3% and 6%, compared to ‘*no management*’ (518 gC m^−2^ y^−1^). The ‘*patch clearcut*’ simulation had similar NPP (509 gC m^−2^ y^−1^) to ‘*no management*’, while the thinning options showed lower values (500 - 508 gC m^−2^ y^−1^) corresponding to a 3% to 4% decrease. All management options showed lower BA values (34 - 38 m^2^ ha^−1^) compared to ‘*no management*’, corresponding to a relative change between −8% (‘*light thinning*’) and −19% (‘*shelterwood B*’). As for pCWS, all options except ‘*natural regeneration*’ (76 tC ha^−1^) had greater values than ‘*no management*’ (121 tC ha^−1^), with the thinning and shelterwood options exhibiting similar values (177 - 193 tC ha^−1^), corresponding to a 45% to 55% increase (Table 3; Figure 3 and 4; Figure S3).

## Discussion

### Model evaluation

The 3D-CMCC-FEM reproduced the development of a Laricio pine stand in the Bonis watershed over a 58 year span. Our evaluation of stand attributes showed that, starting from the establishment of the plantation in 1958, the simulated mean stand DBH and tree density fell reasonably well within the measured range of two independent datasets: average values for low and high density Laricio pine plantations in the area between 1986 and 1999 (Callegari et al.; 2003), and the forest plots surveyed between 1993 and 2016. The model was also able to simulate historical management activities and their effects on forest development. Indeed, the simulation included a thinning of 25% of stand BA that took place in 1993 at the stand, which was reflected by the reduction in tree density in that year and a slight increase in the growth rate of mean stand DBH in the following years (0.6 cm y^−1^ after the thinning vs. 0.3 cm y^−1^ before the thinning).

Furthermore, the model was able to reproduce the mean seasonal cycle of daily GPP as obtained by the eddy covariance tower with sufficient accuracy, supporting previous assessments of model performance (Collalti et al., 2014, 2016, 2018; Alessio Collalti et al., 2020; Dalmonech et al., 2022; Engel et al., 2021; Marconi et al., 2017). The R^2^ of 0.69 is in line with previous evaluations of simulated daily GPP across northern European forest sites (average R^2^ across three sites = 0.73; Collalti et al., 2018), while the ME of 0.61 is within the range found for daily GPP simulated with other process-based models (0.42 - 0.84 in Bagnara et al., 2015; 0.61 - 0.98 in Minunno et al., 2016).

### Impacts of climate change

In the first half of the XXI century, both RCPs projected similar increments in mean annual temperature and vapor pressure deficit with no significant changes in the amount of precipitation for the Bonis watershed. These trends were mirrored by a positive tendency for all output variables in the NF time window. The GPP, NPP, pCWS and BA of the simulated Laricio pine stand seemingly benefitted from the fertilizing effect of increased atmospheric CO_2_ concentration, the lengthening of the growing season and sufficient water availability (Gea-Izquierdo et al., 2017; Kramer et al., 2000; Simioni et al., 2020). Conversely, in the second half of the XXI century, a reduction in precipitation and an increase in temperature – in line with previous estimates for the Mediterranean basin (Lionello and Scarascia, 2018; Santini et al., 2014) – were leading to a decrease of all the variables with the exception of GPP. These changes were more pronounced under the most emission-intensive scenario and toward the end of the century, negatively affecting the ability of Laricio pine stands to absorb and to store C. Indeed, despite a very modest increase in GPP, our simulations predicted a moderate decrease of pCWS and BA, and a strong decrease in NPP of Laricio pine stands. These changes affected all management options regardless of the climate scenario. The decline in water availability is likely responsible for an increased water stress, which could offset the positive effects of increased atmospheric CO_2_ concentrations on photosynthesis (Cinnirella et al., 2002), while higher temperatures favor autotrophic respiration and photorespiration (Dusenge et al., 2019; Gea-Izquierdo et al., 2017; Lindner et al., 2010). If autotrophic respiration increases more than GPP, then NPP decreases proportionally and C-stocks and BA increase at a slower rate (Alessio Collalti et al., 2020). Previous studies already highlighted the negative effect of temperature and soil moisture scarcity on leaf development and tree growth for forests in general and, more in particular, for Laricio pines (Cinnirella et al., 2002; Mazza et al., 2018). However, the emergence of pervasive acclimation mechanisms (e.g., changes in C-allocation for reserve accumulation) in this species could reduce forest vulnerability to extreme events, thus preventing extensive dieback episodes (Cinnirella et al., 2002; Mazza et al., 2018). Nonetheless, indirect effects of climate change, including increased vulnerability of trees to pathogen attacks, could lead to higher mortality rates in spite of physiological adaptations (Gentilesca et al., 2017; Resco De Dios et al., 2007). Recent studies have shown the ambiguity in the responses of forests to both warming and enriched atmospheric CO_2_ concentration (Rezaie et al., 2018), probably related to site-specific factors (e.g. forest age, forest structure, soil nutrient availability and microclimate). While Central and Northern Europe seem to show a general increase in both C-sequestration and C-stocks in the short-to medium-term (Reyer et al., 2015), the impact of increasing droughts and disturbance risk will likely outweigh positive trends in Southern Europe, with an expected decline in the productivity of the Mediterranean region (Lindner et al., 2010; Reyer et al., 2014; Simioni et al., 2020). In this respect, the Bonis experimental watershed represents a unique experimental site with mountain climate at the center of the Mediterranean basin. These features make it particularly exposed to the effects of climate change, hence its likely role of sentinel of future changes in forest dynamics for the whole region.

### Impacts of forest management

Regardless of the short- to long-term reductions in C-fluxes and C-stocks due to increased temperature and lower precipitation, the effect of management on forest attributes largely outplays that of climate change, in line with previous findings for Mediterranean pine forests (del Río et al., 2017) and other European forests (e.g., Akujärvi et al., 2019; Gutsch et al., 2018). Therefore, the choice of far-sighted management options is key to the future of Laricio pine stands in the Bonis watershed, with the aim of preserving and enhancing primary production and carbon storage capacity over time, improving forests resilience to biotic and abiotic stresses, as well as promoting their structural complexity and the multiple ecosystem functions (Scarascia-Mugnozza et al., 2000). The present study aimed at narrowing the knowledge gap about the potential benefits of alternative forest management options for pine plantations under climate change, which is of paramount importance in areas close to the geographical limit of the distribution of pine species like the Bonis watershed (Navarro-Cerrillo et al., 2019).

Our simulations showed that, in the first half of the XXI century, the lack of management interventions led to higher C-fluxes (i.e., GPP and NPP) and BA, as opposed to production-oriented management strategies involving clear-cutting or the shelterwood system, which abruptly slowed down C-fluxes because of the strong reduction in leaf area and in situ standing biomass. Yet, such commercial forest-oriented options showed to maximize C-fluxes in the second half of the XXI century as a response to regeneration or replanting. Despite these fluctuations, the overall effect on C-fluxes of different management options over the 2006 - 2095 period was modest, with a relative change range between −6% and +7% compared to ‘*no management*’. These results might allude that either forest management is counterbalancing the apparently positive effects of warming and increasing atmospheric CO2 concentration, or that the Laricio pine has already reached its suitability optimum for this particular geographic area. However, it has been previously demonstrated that the lack of forest management in pine plantations might increase inter-tree competition, hence vulnerability to drought stress (Manrique-Alba et al., 2020; Martín-Benito et al., 2010; Navarro-Cerrillo et al., 2019).

Furthermore, unmanaged pine plantations of the Mediterranean basin are simplified ecosystems composed of high-density, even-aged stands with arrested succession and at risk of catastrophic events like wildfires and pests outbreaks (Ruiz-Benito et al., 2012; Scarascia-Mugnozza et al., 2000). In this study we simulated a ‘*natural regeneration*’ option in which the unmanaged standing biomass is eliminated after a simulated destructive event (i.e., a wildfire). While the average C-sink is similar to the other management options, all the on-site C returns to the atmosphere as an effect of the simulated disturbance. As this scenario represents an increasingly likely outcome in Mediterranean pine plantations under climate change, forest managers should prioritize active management options aimed at reducing fire risk by decreasing the fuel load. Among these options, thinning interventions are particularly promising, as they have demonstrated to reduce fireline intensity while avoiding emissions from prescribed burning (Rabin et al., 2022). The simulated thinning options exhibited minor reductions in C-fluxes and BA, compared to the absence of management, along the whole simulation. The ‘*light thinning*’ option (28% reduction of BA every 15 years), in particular, showed the lowest decrease of the above mentioned variables among all active management strategies. Conversely, pCWS were maximized under the ‘*heavy thinning*’ and ‘*shelterwood B*’ options, which involved the strongest removals of BA (35.5% reduction of BA every 15 years and 28.5% of BA every 10 years, respectively).

Previous studies highlighted the role of management strategies comprising a reduction of tree density (i.e., thinning and shelterwood) in improving overall forest health in the Mediterranean region (Brichta et al., 2020; del Río et al., 2017; Manrique-Alba et al., 2020; Martín-Benito et al., 2010; Navarro-Cerrillo et al., 2019; Prévosto et al., 2011; Ruiz-Benito et al., 2012). In the shelterwood system, stand density is reduced to increase light availability, with positive effects on the growth of naturally established seedlings (Prévosto et al., 2011). Shelterwood regeneration of pine species was found to be more favorable with respect to microsite characteristics and of greater quality compared to replanting after clear-cut – especially after a heavy reduction of initial stand density – making it a potentially useful management option to mitigate the negative effects of climate change (Brichta et al., 2020). Similarly, thinning interventions have been observed to reduce competition for water, light and soil nutrients – thus increasing photosynthetic rates – as well as improving both carbon and water use-efficiency and C-uptake capacity of remaining trees (Collalti et al., 2020; Manrique-Alba et al., 2020; Martín-Benito et al., 2010; Navarro-Cerrillo et al., 2019; Rezaie et al., 2018). In particular, moderate to heavy thinning interventions (between 25 to 50% reduction of stand BA) have been recommended as a drought adaptation measure for Mediterranean pine forests with long-lasting positive effects (Manrique-Alba et al., 2020). Furthermore, heavy thinning was found to increase the C-sequestration potential of these environments by compensating the loss of on-site C with an increased total C-stock when harvested woody products are taken into account (del Río et al., 2017). Our results for a Laricio pine stand at the Bonis watershed were consistent to the above mentioned findings. Thinnings represent a viable management option for the study area that maximizes the potential C-stocks while providing improved conditions in relation to secondary climate change effects. On the other hand, the shelterwood options represent a halfway alternative between patch clearcut and thinnings, that can be used to renaturalize Laricio pine forests, with cascading positive effects on the local water balance and hydrogeological risk reduction.

### Assumptions and caveats

The 3D-CMCC-FEM allowed to simulate several management options for Laricio pine plantations at Bonis watershed under different climate scenarios considering biogeochemical, biophysical, physiological and stand development processes. In the current version, the model was unable to simulate some forest disturbances that are likely to impact our study area like recurrent wildfires and pest outbreaks. However, we explicitly simulated a single destructive event under the ‘*natural regeneration*’ option, consisting in the complete removal of the standing biomass after 80 years from the planting, followed by natural regeneration. Although limited in scope, such simulation provides an overview of the effects of perturbations that might potentially occur to Laricio pine plantations in the absence of proactive management in the area. We also recognize that more management options than the ones we simulated are available. Yet, our scenarios cover several objectives including post-disturbance management, wood production and renaturalization and reflect the state-of-the-science of management types applied to this region of the Italian Apennines (Cantiani et al., 2018). Furthermore, the model does not account for the effect of soil nutrients on tree growth. Yet, nutrient availability is generally considered a secondary driver of tree growth in Laricio pine forests, which are however mainly limited by soil moisture (Mazza et al., 2018). Finally, the simulations did not include species replacement due to competition and colonization. However, the forests at the Bonis watershed are dominated by Laricio pines, both natural and artificial, which are likely to recolonize gaps in the absence of proactive replanting of other tree species.

## Conclusions

Overall, our 137-year simulation showed that climate change will affect the development of Laricio pine plantations at the Bonis watershed, with profound impacts on C-sinks and C-stocks especially in the second half of the century. However, the choice of future management will exert an even stronger effect on the C-sink and C-stock capacity of such forests. Therefore, planning appropriate management options aimed at maintaining and enhancing these features, while favoring the renaturalization of these environments, is key to allow the future provision of forest ecosystem services in the area. Among the investigated options, thinning interventions represent the most promising management practice, also considering their documented contribution to increasing drought resistance and reducing fire risk. The present work provided a first overview of the joint effect of climate change and management on one of the southernmost European forest sites, with direct implications for the planning of adaptive management strategies in Mediterranean pine forests. Yet, further studies are required to assess the impact of recurrent stand disturbances, changes in soil nutrient concentrations and species replacement on multiple ecosystem services.

## Acknowledgements

This study presented the results obtained within the ALForLab project (PON03PE_00024_1) co-funded by the National Operational Program for Research and Competitiveness (PON R&C) 2007–2013, through the European Regional Development Fund (ERDF) and national resources (Revolving Fund - Cohesion Action Plan PAC). RT has been supported by the Italian Ministry of University and Research (FOE-2019) under the project ‘Climate Changes’ (CNR DTA. AD003.474.029) and by LifeWatch Italy through the project LifeWatchPLUS (CIR-01_00028). DD acknowledges funding by the project OT4CLIMA which was funded by the Italian Ministry of Education, University and Research (D.D. 2261 del 6.9.2018, PON R&I 2014-2020 e FSC). The authors would like to thank G. Scarascia-Mugnozza and G. Pellicone and colleagues at ISAFOM-CNR-RENDE for providing us some of the field data used in this work. The 3D-CMCC-FEM model code is publicly available and can be found on the GitHub platform at: https://github.com/Forest-Modelling-Lab/3D-CMCC-FEM). All data supporting this study are publicly available at XXX. Requests for additional material should be addressed to the corresponding author.

## Author contributions

**Riccardo Testolin**: Conceptualization, Data curation, Formal analysis, Investigation, Methodology, Resources, Software, Validation, Visualization, Writing - original draft. **Maurizio Bagnara**: Resources, Writing - review & editing. **Daniela Dalmonech**: Conceptualization, Methodology, Resources, Software, Supervision, Writing - original draft. **Ettore D’Andrea**: Writing - review & editing. **Gina Marano**: Conceptualization, Methodology, Resources, Software, Writing - original draft. **Giorgio Matteucci**: Writing - review & editing. **Sergio Noce**: Resources, Writing - review & editing. **Alessio Collalti**: Conceptualization, Funding acquisition, Methodology, Resources, Software, Supervision, Project administration, Writing - original draft.

## Notes

### Competing Interest Statement

The authors have declared no competing interest.

