## Supplementary information for "Simulating alternative forest management in a changing climate on a *Pinus nigra* subsp. *laricio* plantation in Southern Italy"

**Table S1.** Parameterization file for 3D-CMCC-FEM used in the present study for *Pinus nigra* subsp. *laricio.*

| Parameter | Value | Species | Source | Description |
|---|---|---|---|---|
| **LIGHT_TOL** | 4 | *P. nigra* | Patenaude et al., 2008 | Light tolerance: 4 = very shade intolerant (cc = 90%), 3 = shade intolerant (cc = 100%), 2 = shade tolerant (cc = 110%), 1 = very shade tolerant (cc = 120%) (dimensionless) (cc = canopy cover) |
| **PHENOLOGY** | 1.2 | *P. nigra* | | Phenology of species: 0.1 = deciduous broadleaf, 0.2 = deciduous needleleaf, 1.1 = broadleaf evergreen, 1.2 = needleleaf evergreen (dimensionless) |
| **K** | 0.5 | *P. nigra* | Navarro-Cerrillo et al., 2016 | Extinction coefficient for absorption of PAR by canopy (ratio) |
| **ALBEDO** | 0.126 | *P. nigra* | Moricz et al., 2018 | Canopy albedo (ratio) |
| **SLA_AVG0** | 5 | *P. nigra* | Navarro-Cerrillo et al., 2016 | Average specific leaf area (juvenile) ($m^2Kg\ DM^{-1}$) |
| **SLA_AVG1** | 4 | *P. nigra* | Navarro-Cerrillo et al., 2016 | Average specific leaf area (mature) ($m^2Kg\ DM^{-1}$) |
| **TSLA** | 4 | *P. nigra* | Navarro Cerrillo et al., 2016 | Age at which SLA_AVG = (SLA_AVG0 + SLA_AVG1) / 2 (yr) |
| **SLA_RATIO** | 2.52 | *P. pinaster* | Mollicone et al. 2002 | Ratio of shaded to sunlit projected SLA (ratio) |
| **LAI_RATIO** | 2.6 | *P. pinaster* | Chiesi et al., 2007 | Ratio of all-sided to projected leaf area (ratio) |
| **FRACBB0** | 0.5 | *P. nigra* | Navarro Cerrillo et al., 2016 | Branch and bark fraction (juvenile) (ratio) |
| **FRACBB1** | 0.1 | *P. nigra* | Navarro Cerrillo et al., 2016 | Branch and bark fraction (mature) (ratio) |
| **TBB** | 5 | *P. nigra* | Navarro Cerrillo et al., 2016 | Age at which FRACBB = (FRACBB0 + FRACBB1 ) / 2 (yr) |
| **RHO0** | 0.43 | *P. nigra laricio* | Patenaude et al., 2008 | Minimum basic density (juvenile) ($Mg\ DM\ /m^{-3}$) |

| | | | | |
|---|---|---|---|---|
| **RHO1** | 0.43 | *P. nigra laricio* | Patenaude et al., 2008 | Minimum basic density (juvenile) (Mg DM $m^{-3}$) |
| **TRHO** | 4 | *P. nigra* | Navarro-Cerrillo et al., 2016 | Age at which RHO = (RHO0 + RHO1) / 2 (yr) |
| **COEFFCOND** | 0.05 | *P. nigra* | Navarro-Cerrillo et al., 2016 | Controls stomatal response to VPD (mbar) |
| **BLCOND** | 0.2 | *P. nigra* | Navarro-Cerrillo et al., 2016 | Canopy boundary layer conductance (m $s^{-1}$) |
| **MAXCOND** | 0.0021 | *P. nigra laricio* | Estimated from Lapa et al., 2017 | Maximum leaf (stomatal) conductance (m $s^{-1}$) |
| **CUTCOND** | 2.5E–05 | *P. pinaster* | Mollicone et al., 2002 | Cuticular conductance (m $s^{-1}$) |
| **MAXAGE** | 400 | *P. nigra* | Grossoni, 2014 | Controls rate of physiological decline of forest (yr) |
| **RAGE** | 0.95 | *P. nigra* | Navarro-Cerrillo et al., 2016 | Relative age to give f(AGE) = 0.5 (dimensionless) |
| **NAGE** | 4 | *P. nigra* | Navarro-Cerrillo et al., 2016 | Power of relative age in f(AGE) (dimensionless) |
| **GROWTHTMIN** | 0 | *P. nigra* | Navarro-Cerrillo et al., 2016 | Minimum temperature for growth (°C) |
| **GROWTHTMAX** | 35 | *P. nigra* | Navarro-Cerrillo et al., 2016 | Maximum temperature for growth (°C) |
| **GROWTHTOPT** | 15 | *P. nigra* | Navarro-Cerrillo et al., 2016 | Optimum temperature for growth (°C) |
| **SWPOPEN** | –0.4 | *P. nigra laricio* | Leborgeois et al., 1998 | Leaf water potential (start of reduction) (MPa) |
| **SWPCLOSE** | –1.7 | *P. nigra laricio* | Leborgeois et al., 1998 | Leaf water potential (end of reduction) (Mpa) |
| **OMEGA** | 0.5 | Needle-leaf | Arora et al., 2005 | Controls sensibility of allocation to changes in water and light availability (dimensionless) |
| **S0** | 0.6 | *Pinus sp.* | Deduced from Dewar et al., 1994 | Stem allocation factor (ratio) |
| **R0** | 0.2 | *Pinus sp.* | Deduced from Dewar et al., 1994 | Root allocation factor (ratio) |
| **F0** | 0.2 | *Pinus sp.* | Deduced from Dewar et al., 1994 | Foliage allocation factor (ratio) |
| **FRUIT_PERC** | 0.2 | *P. sylvestris* | Yuste et al., 2005 | Fraction of NPP allocated for reproduction during the prescribed seasonal period (ratio) |

| | | | | |
|---|---|---|---|---|
| **FINE_ROOT_LEAF** | 0.7 | *P. pinaster* | Mollicone et al. 2002 | Fine root C:leaf C (ratio) |
| **COARSE_ROOT_STEM** | 0.16 | *P. pinaster* | Mollicone et al. 2002 | Coarse root C:stem C (ratio) |
| **LIVE_TOTAL_WOOD** | 0.071 | *P. pinaster* | Mollicone et al. 2002 | Live C:total wood C (ratio) |
| **N_RUBISCO** | 0.04 | *P. pinaster* | Chiesi et al., 2007 | Fraction of leaf N in Rubisco (ratio) |
| **CN_LEAVES** | 42 | *P. pinaster* | Chiesi et al., 2007 | C:N of leaves (ratio) |
| **CN_FINE_ROOTS** | 58 | *P. pinaster* | Chiesi et al., 2007 | C:N of fine roots (ratio) |
| **CN_LIVEWOOD** | 58 | *P. pinaster* | Chiesi et al., 2007 | C:N of live woods (ratio) |
| **LEAF_FINEROOT_TURNOVER** | 0.26 | *P. nigra* | Withington et al., 2006 (1/3.84 = only leaf, fine root would be 0.6) | Average annual leaf and fine root turnover ($yr^{-1}$) |
| **LIVEWOOD_TURNOVER** | 0.05 | FIXED | Poulter et al., 2010 | Annual livewood turnover ($yr^{-1}$) |
| **SAPWOOD_TURNOVER** | 0.05 | FIXED | Poulter et al., 2010 | Annual sapwood turnover ($yr^{-1}$) |
| **DBHDCMIN** | 0.14 | *P. sylvestris* | Collalti et al., 2019 | Minimum DBH to crown diameter (ratio) |
| **SAP_A** | 0.32 | *P. pinaster* | Deduced from Delzon et al., 2004 | Scaling coefficient in sapwood area to DBH relationship (dimensionless) |
| **SAP_B** | 1.9 | *P. pinaster* | Deduced from Delzon et al., 2004 | Scaling coefficient in sapwood area to DBH relationship (exp) (dimensionless) |
| **SAP_LEAF** | 1500 | *P. nigra* | Margolis et al., 1995 | Leaf area to sapwood area (ratio) |
| **SAP_WRES** | 0.05 | Evergreen | Schwalm & Ek, 2004; Collalti et al., 2020 | Sapwood to reserve biomass: 0.11 = deciduous, 0.05 = evergreen (ratio) |
| **STEMCONST_P** | 0.043808 | *P. nigra* | Navarro-Cerrillo et al., 2016 | Scaling coefficient in stem mass to DBH relationship (dimensionless) |
| **STEMPOWER_P** | 2.4975 | *P. nigra* | Navarro-Cerrillo et al., 2016 | Scaling coefficient in stem mass to DBH relationship (exp) (dimensionless) |

| | | | | |
|---|---|---|---|---|
| **CRA** | 22.93 | *P. nigra laricio* | Estimated from plot data | Chapman-Richards asymptotic maximum height (m) |
| **CRB** | 0.15 | *P. nigra laricio* | Estimated from plot data | Chapman-Richards exponential decay parameter (dimensionless) |
| **CRC** | 2.64 | *P. nigra laricio* | Estimated from plot data | Chapman-Richards shape parameter (dimensionless) |
| **CROWN_A** | 0.4 | *P. nigra laricio* | Estimated from plot data | Scaling coefficient in crown length to height relationship (dimensionless) |
| **CROWN_B** | 0.99 | *P. nigra laricio* | Estimated from plot data | Scaling coefficient in crown length to height relationship (exp) (dimensionless) |
| **SEXAGE** | 30 | *P. nigra* | Van Haverbeke, 1990 | Age at sexual maturity (yr) |

**Table S2.** Mean of climatic variables for three climate change scenarios within the NF and FF time windows. Relative changes compared to the baseline NOCC scenario are reported in brackets.

| | **Near future (2025 - 2055)** | | | **Far future (2065 - 2095)** | | |
|---|---|---|---|---|---|---|
| | **NOCC (baseline)** | **RCP4.5** | **RCP8.5** | **NOCC (baseline)** | **RCP4.5** | **RCP8.5** |
| **Vapor pressure deficit (mbar)** | 7.4 | 8.2 (13) | 8.6 (18) | 7.3 | 9.4 (31) | 11.3 (59) |
| **Precipitation (mm)** | 907 | 921 (6) | 794 (–8) | 965 | 753 (–20) | 736 (–22) |
| **Mean temperature (°C)** | 13.2 | 14.4 (9) | 15.0 (14) | 12.9 | 15.9 (23) | 17.9 (39) |
| **Atmospheric $CO_2$ (ppmv)** | 365 | 461 (26) | 494 (35) | 364 | 530 (45) | 761 (109) |

**Table S3.** Mean of relative change in simulation outputs for the RCP4.5 and RCP8.5 scenarios compared to NOCC within the NF and FF time windows. The values have been averaged across all management options.

| | Near future (2025 - 2055) | | Far future (2065 - 2095) | |
|---|---|---|---|---|
| | **RCP4.5** | **RCP8.5** | **RCP4.5** | **RCP8.5** |
| **GPP** | 4.7 | 7.8 | 3.0 | 1.9 |
| **NPP** | 0.4 | 1.3 | –11.7 | –23.0 |
| **pCWS** | 0 | 0.7 | –1.8 | –2.2 |
| **BA** | 1.5 | 2.3 | –1.2 | –1.9 |

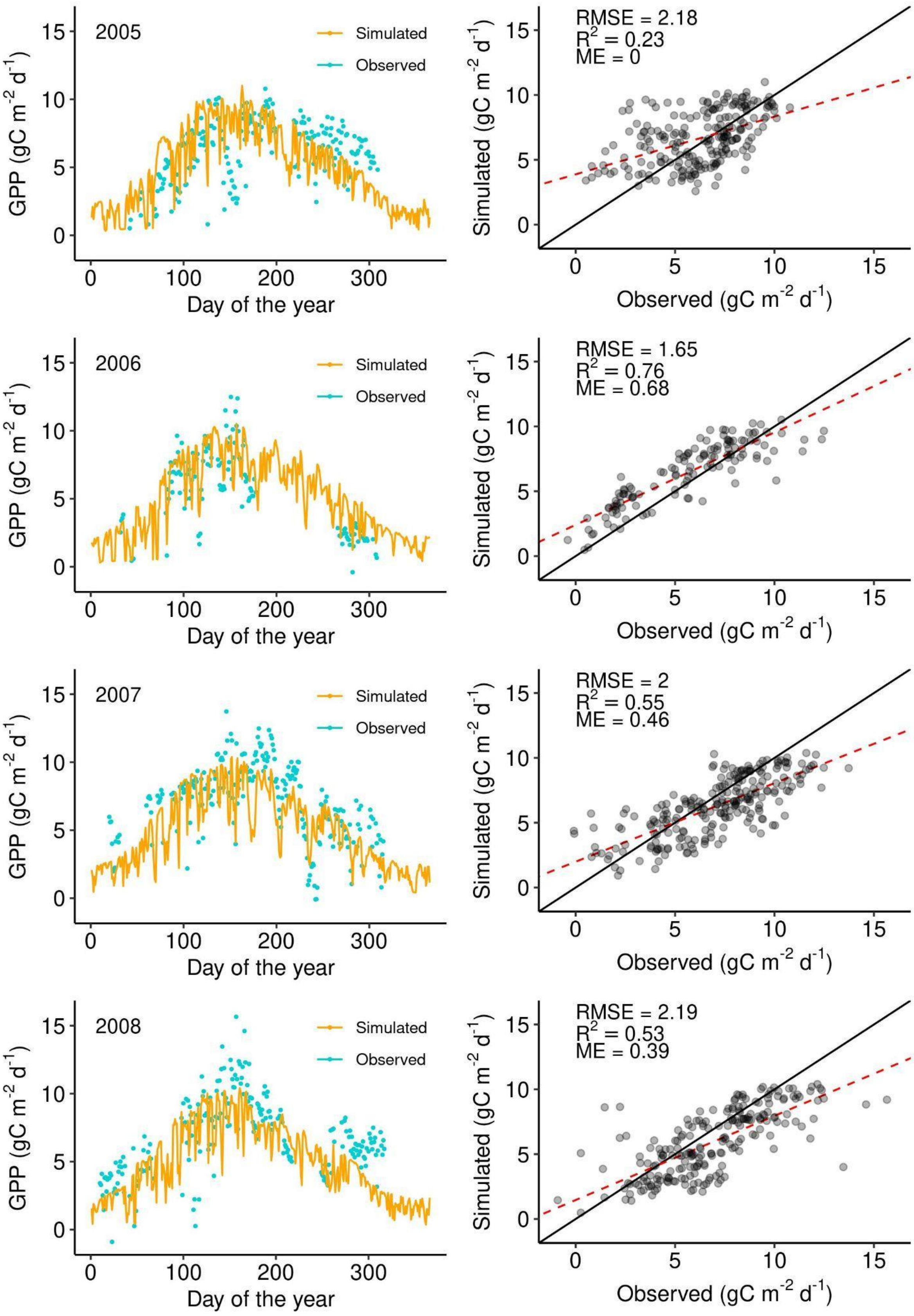


**Figure S1.** Evaluation of simulated daily GPP using the 3D-CMCC-FEM for the years 2005 - 2008 against the values measured by the eddy covariance tower at the Bonis watershed.

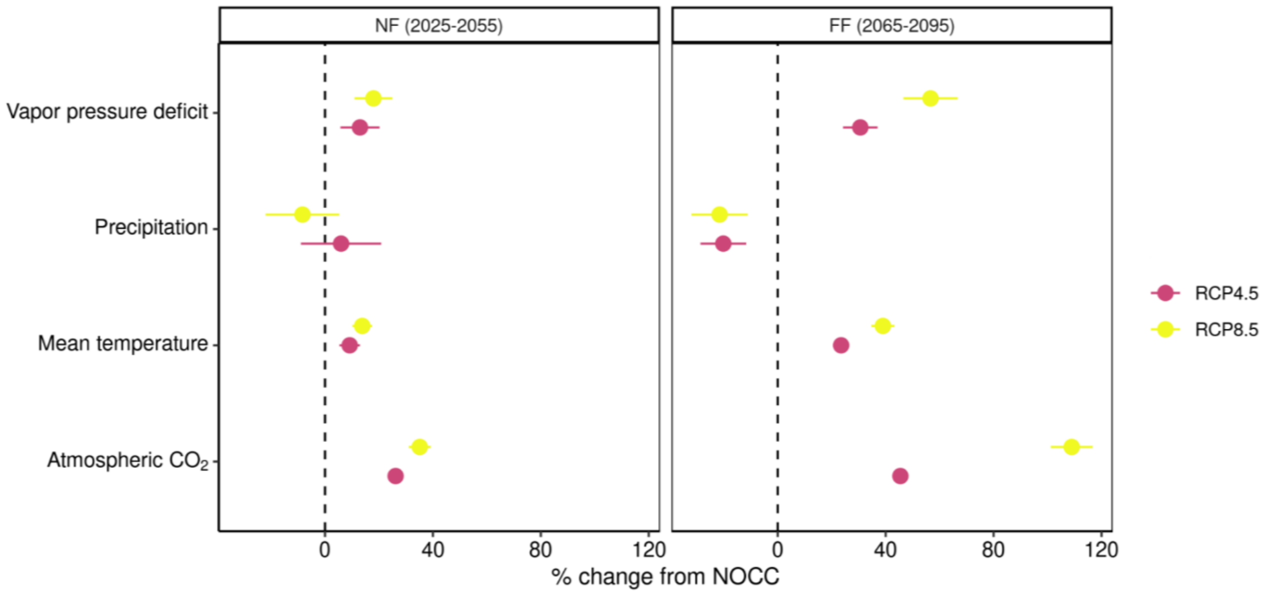


**Figure S2.** Relative change of climatic variables according to RCP4.5 and RCP8.5 climate scenarios compared to the baseline NOCC scenario within the NF and FF time windows. The dots are the mean percentage change calculated within each time window. The error bars are the 95% confidence intervals.

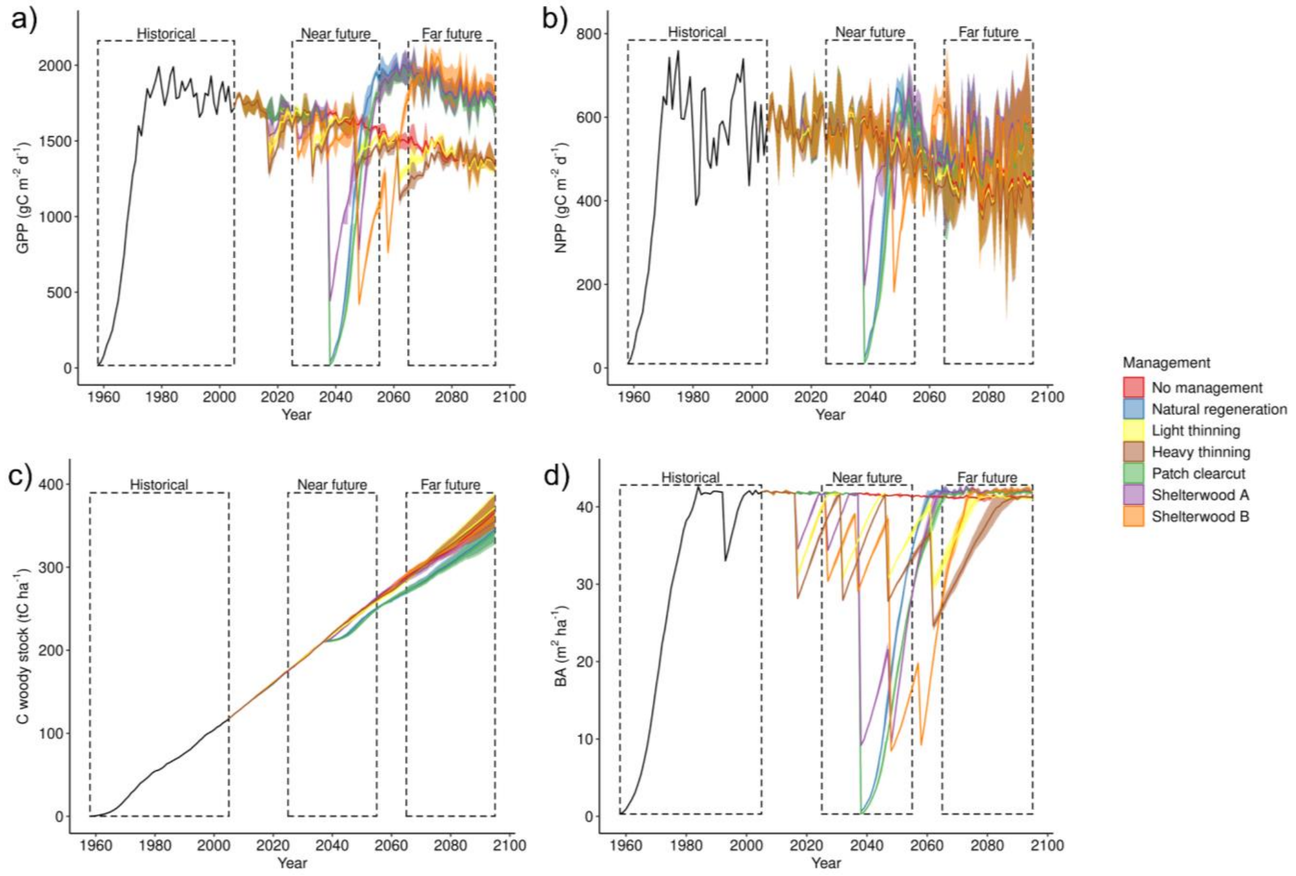


**Figure S3.** Simulated GPP (a), NPP (b), pCWS (c) and BA (d) according to seven management options. Black lines are the historical simulations from 1958 to 2005. Solid lines from 2006 onwards are the mean outputs produced by different climate scenarios (NOCC, RCP4.5, RCP8.5) for each management option. Shaded areas are the interval between the maximum and minimum values for each management option.
